# Temporal and spatial characterization of HIV/SIV infection at anorectal mucosa using rhesus macaque rectal challenge model

**DOI:** 10.1101/2023.02.22.529624

**Authors:** Danijela Maric, Lisette Corbin, Natalie Greco, Ramon Lorenzo-Redondo, Michael D. McRaven, Ronald S. Veazey, Thomas J. Hope

**Affiliations:** Northwestern University Feinberg School of Medicine, Department of Cell and Developmental Biology, Chicago, Illinois, USA; Emory University School of Medicine, Atlanta, Georgia, USA; Parker Institute for Cancer Immunotherapy, San Francisco, California, USA; Northwestern University Feinberg School of Medicine, Department of Medicine, Division of Infectious Diseases, Chicago, Illinois, USA; Center for Pathogen Genomics and Microbial Evolution, Northwestern University Havey Institute for Global Health, Chicago, Illinois, USA; Tulane National Primate Research Center, Division of Comparative Pathology, Covington, Louisiana, USA

**Keywords:** Th17 T cells, iDCs, HIV/SIV, receptive anal intercourse, mucosal tissues, SIVmac239, rhesus macaque

## Abstract

The study described herein is a continuation of our work in which we developed a methodology to identify small foci of transduced cells following rectal challenge of rhesus macaques with a non-replicative luciferase reporter virus. In the current study, the wild-type virus was added to the inoculation mix and twelve rhesus macaques were necropsied 2-4 days after the rectal challenge to study the changes in infected cell phenotype as the infection progressed. Relying on luciferase reporter we noted that both anus and rectum tissues are susceptible to the virus as early as 48h after the challenge. Small regions of the tissue containing luciferase-positive foci were further analyzed microscopically and were found to also contain cells infected by wild-type virus. Phenotypic analysis of the Env and Gag positive cells in these tissues revealed the virus can infect diverse cell populations, including but not limited to Th17 T cells, non Th17 T cells, immature dendritic cells, and myeloid-like cells. The proportions of the infected cell types, however, did not vary much during the first four days of infection when anus and rectum tissues were examined together. Nonetheless, when the same data was analyzed on a tissue-specific basis, we found significant changes in infected cell phenotypes over the course of infection. For anal tissue, a statistically significant increase in infection was observed for Th17 T cells and myeloid-like cells, while in the rectum, the non-Th17 T cells showed the biggest temporal increase, also of statistical significance.

**IMPORTANCE:** Men who have sex with men are at the highest risk of acquiring HIV via receptive anal intercourse. Understanding what sites are permissive to the virus, and what the early cellular targets are is critical for development of effective prevention strategies to control HIV acquisition amid receptive anal intercourse. Our work sheds light on the early HIV/SIV transmission events at the rectal mucosa by identifying the infected cells and highlights the distinct roles that different tissues play in virus acquisition and control.

## INTRODUCTION

HIV transmission via receptive anal intercourse (RAI) continues to be a major driver in the HIV epidemic [1-3]. While both men and women are susceptible to acquiring the virus via this route, men who have sex with men (MSM) are at the highest risk, representing nearly 70% of all new cases in the USA [4-6]. While many studies have focused on studying transmission at the permissive single-layer columnar tissues of the rectum [5, 7-9], the more robust multilayered squamous epithelia of the anus tissues have for the most part been ignored. Several groups have attempted to gain insights into early cellular targets of HIV using tissue digestion and flow cytometry [10, 11]; however, it is known that this method can result in losses in certain cell populations due to mechanical and enzymatic digestions [12-14]. Lastly, the majority of the work involving transmission via rectal route was conducted several days after the initial challenge [15-17] hence little information is available regarding the transmission events in the earliest days after viral exposure. To develop effective prevention strategies, it is critical to understand what sites are permissive to the virus and what the early cellular targets at these sites are.

In our previous work aimed at better understanding HIV transmission during rectal/anal intercourse (RAI), rhesus macaques were intrarectally challenged with a single-round SIV-based dual reporter vector that expresses luciferase [18, 19] and near-infrared fluorescent protein 670 (iRFP670) [20, 21] upon transduction of target cells [22]. Regions of tissue containing luminescent transduced cells were identified macroscopically using an *in vivo* imaging system, and individual transduced cells expressing fluorescent protein iRFP670 were identified and characterized microscopically [22]. This system revealed that anal and rectal tissues are both susceptible to transduction just 48h after rectal challenge [22]. Detailed phenotypic analysis revealed that most of the transduced cells are CCR6-positive T cells—the vast majority of which expressed RORγT, a Th17 lineage-specific transcription factor [22-26]. Interestingly, the second most common target was not another T cell subtype but rather immature dendritic cells (iDCs) [22]. Both cell types occur at a low frequency in the anorectal tissue, yet had higher rates of infection, indicating that they are the preferential initial targets of HIV/SIV in rectal transmission [22]. We have previously reported that Th17 lineage cells and iDCs are also the preferential targets of SIV during vaginal transmission [27].

One downside of using a single-round, replication-incompetent virus is that it does not allow for the study of virus-host cell responses to infection nor changes in the phenotype of infected cells over time. To overcome these limitations and contribute to the outstanding questions in the field, we utilized an inoculum containing the mixture of the luciferase reporter virus and wild-type SIVmac239, an approach successfully used to identify the infected cells in female reproductive tract (FRT) after vaginal challenge [27]. By homing in on the sites that demonstrated luciferase expression we were able to identify the cells infected by the wild-type virus and to assess the cellular targets of the wild-type virus in the early days of infection in anorectal mucosa.

Much like in our original study utilizing reporter virus alone [22], we found that anus and rectum are also susceptible to the wild-type virus. The infected cell foci were detectable as early as 48h after rectal inoculation and these infected cell foci grew significantly in size as the infection progressed. By phenotyping infected cells 48-, 72- and 96h post challenge, we found that the infected cell proportions remained relatively constant over the first four days of infection when anorectal tissue was examined as a whole. However, when infected cell phenotypes were compared between tissue types, we observed differences between the two tissue types. Most notably, for anal tissue, the biggest temporal change occurs in infection rates of Th17 T cells and myeloid-like cells, both which show significant increases in infection rates from 48-96h. In the rectum, however, the biggest temporal change occurs in infection rates of other T cells, which likewise show a significant increase as infection progressed. This study identifies early cellular targets of HIV/SIV in the anorectal mucosa and highlights the important roles that different tissues play in virus susceptibility and infection progression.

## RESULTS

### Luciferase reporter can be used to find cells infected by the wild-type SIVmac239 in anorectal tissues

In our previous work in the FRT, we used a mixed inoculum including the non-replicating luciferase virus and wild-type SIVmac239 to identify the sites that are susceptible to the virus and to characterize the infected cells 48-hours after vaginal challenge [27]. We showed that non-replicating luciferase virus can be used as a beacon to identify the sites of weakened barrier function and these sites were screened for the presence of the wild-type virus. The presence of infected cells was found throughout the FRT and a close correlation was observed between the luciferase reporter virus and wild-type SIV [27]. Based on this work, here we sought to investigate temporal changes in infection after rectal challenge. Twelve female RMs received small punch biopsies at random and were inoculated intrarectally with a mixture of non-replicative reporter virus JRFL-LI670 and wild-type SIVmac239. Animals were necropsied either 48-, 72- or 96h later (4 animals per group). This study analyzed the entire anus and the adjoining 4-6cm of the rectum. Isolated tissues were soaked in luciferin and examined by *in vivo* imaging system (IVIS), where luciferase reporter from JRFL-LI670 virus was used to identify the sites that are susceptible to the virus. The number, size and distribution of luciferase positive foci differed between the twelve animals (data not shown), consistent with our previous studies where the same or similar luciferase reporter viruses were used to understand rectal and vaginal transmission 48h after the challenge [18, 22, 27]. We could detect robust luciferase activity even 96h post rectal challenge, which helped us identify the small regions of tissues that likely also contain infected cells (Figure 1A). To validate that luciferase signal could be used as a guide in finding the cells infected by the wild-type virus, the regions of tissue with the strongest luciferase signal were flash frozen, cryosectioned, immunolabeled, and imaged by fluorescence microscopy. Since the wild-type virus used in this study is not fluorescently tagged we sought to identify the infected cells by staining for viral proteins Gag and Env (Figure 1B). Immunofluorescence analyses revealed robust protein expression of both viral proteins in a myriad of cells with clearly defined nuclei. These cells had a focal appearance and were present in both anus and rectum tissues.

**Figure 1:**
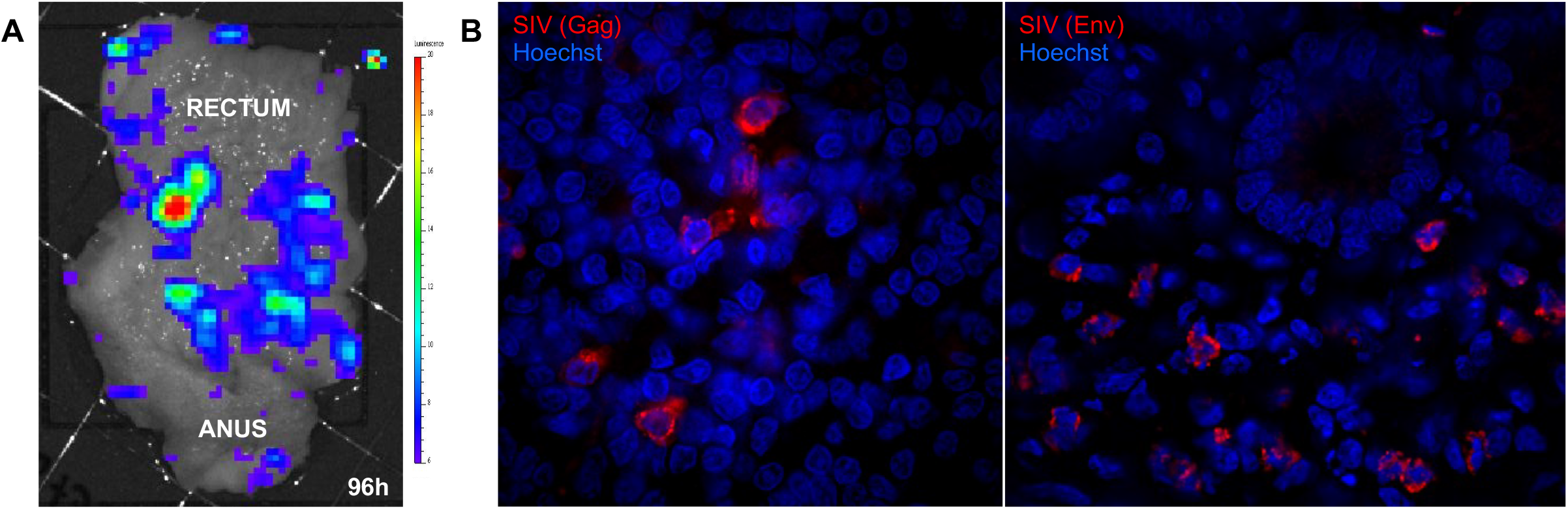
Macroscopic and microscopic identification of SIVmac239 infected cell foci at the anorectal mucosa. Twelve female macaques were rectally inoculated with a mixture of LI67-JRFL virus and wild-type SIVmac239, sacrificed either 48-, 72-or 96h later, and the first 4-6cm distal colon was removed for analysis. (A) Luciferase reporter expression in anorectal tissue 96h after the rectal challenge. Anorectal tissues were cleaned in PBS, soaked in luciferin, and imaged by IVIS to identify the areas of tissue susceptible to the virus. Both anal and rectal poles are indicated. (B) Identification of SIVmac239 infected cells in anorectal tissues. Tissue regions containing high luciferase signals were excised, frozen in OCT blocks, cryosectioned, and stained with SIV Env antibodies KK41 and KK42 (red, Figure 1B left panel) and SIV Gag antibody Ag3.0 (red, Figure 1B right panel). Nuclear dye Hoechst (blue) was used to denote nuclei of the individual cells.

### Infection by SIVmac239 causes expected changes in CD4 and CD3 expression on the infected cells

To further validate infection and to begin to phenotype infected cells, the tissue sections containing infected cells were stained with anti CD4 (HIV-1 receptor) [28, 29] and anti CD3 (T cell marker) [30, 31] antibodies. We noted varying levels of CD4 expression in Env-positive cells. Some infected cells had high CD4 signals, while others had low to no CD4 signal detectable (Figure 2A). Varying CD4 expression in infected cells is consistent with our previously published work in FRT where it was determined that plasma membrane associated CD4 receptors are internalized and degraded as a result of infection [27]. When we looked at CD3 expression we could see that both T cells and non-T cells are susceptible to the virus as early as 48h after rectal challenge (Figure 2B). Moreover, we noticed disparate CD3 expression in some SIV-infected cells compared to uninfected neighboring cells, in agreement with our published work in FRT where we showed that CD3 was internalized as the result of infection (Figure 2C) [27].

**Figure 2:**
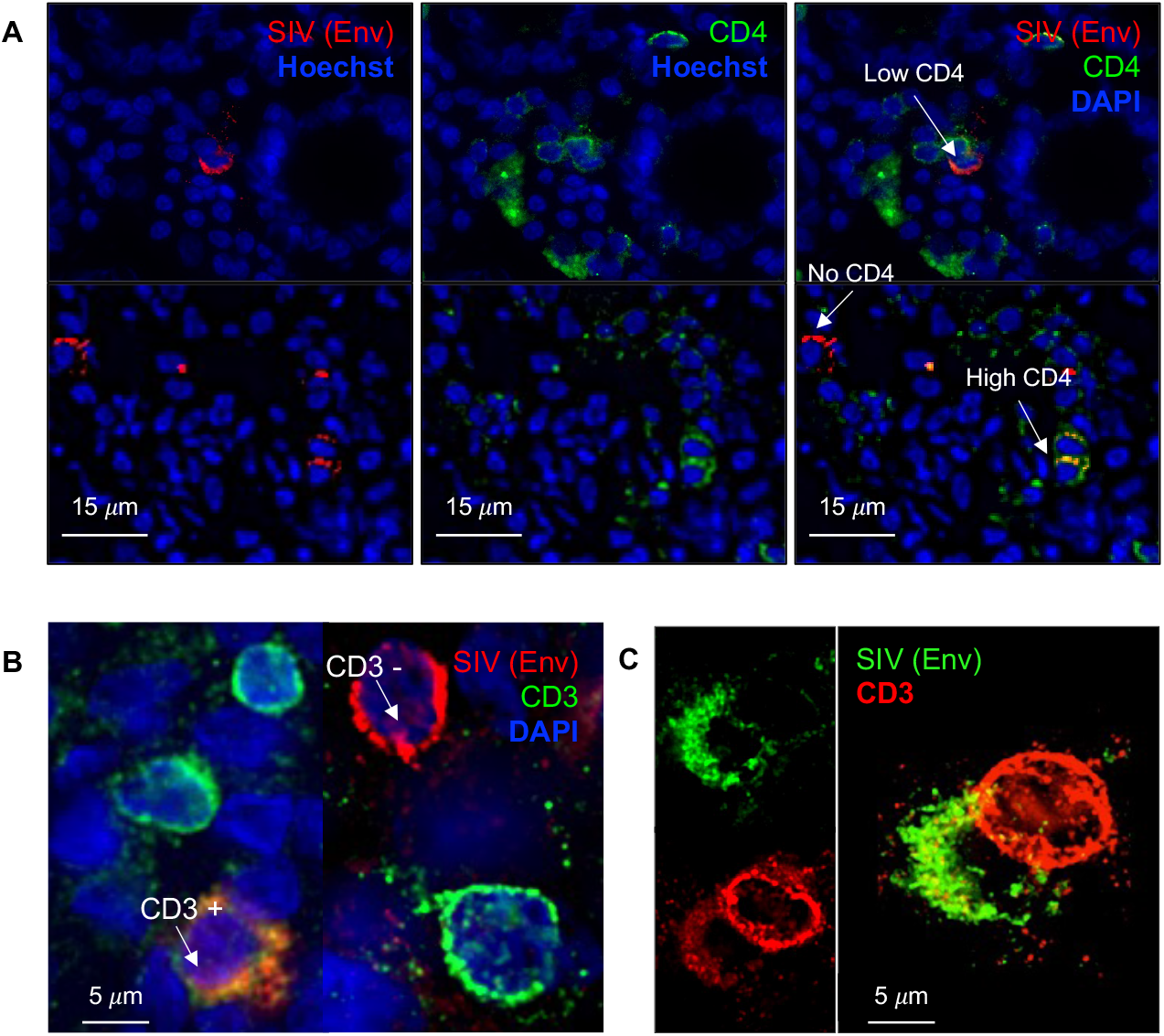
Changes in CD4 and CD3 expression resulting from SIV infection. Cryosections containing SIV-infected cells (red) were immunolabeled with either CD4 or CD3 antibody (green) and nuclear stain Hoechst (blue). (A) each column shows an image of the same field where varying levels of CD4 expression were observed in infected cells. (B) Dual immunostaining with SIV Env antibody and CD3 showed that both T cells and non-T cells are susceptible to the virus. (C) Dual immunostaining with SIV Env antibody and CD3 shows disparate levels of CD3 in some of the infected cells compared to noninfected cells in proximity. *Panel C of this figure was previously shown on the cover of AIDS Research and Human Retroviruses Journal.

### Temporal increase in infected cell foci size in anorectal tissues

To gain insights into time-dependent changes in infected cell distribution and to determine the localization of the infected cells we cryosectioned luciferase-positive tissue blocks and performed immunofluorescence staining for SIV Env protein and examined the tissue sections by fluorescence microscopy. When we compared the size of infected cell foci at 48-, 72- and 96h after rectal challenge we observed pronounced changes in the number of infected cells within individual foci (Figure 3A, anus and Figure 4, rectum). At the 48h time point, we often see individual infected cells dispersed throughout the tissue with small foci usually consisting of only a few of infected cells. By 72h the foci grow notably in size, encompassing several dozen infected cells. Lastly, by the 96h time point the foci grow significantly in size and often stretch across multiple adjacent tissue sections. In both anal and rectal tissue, the infected cells were predominantly localized in the lamina propria with varying distances from the basement membrane.

**Figure 3:**
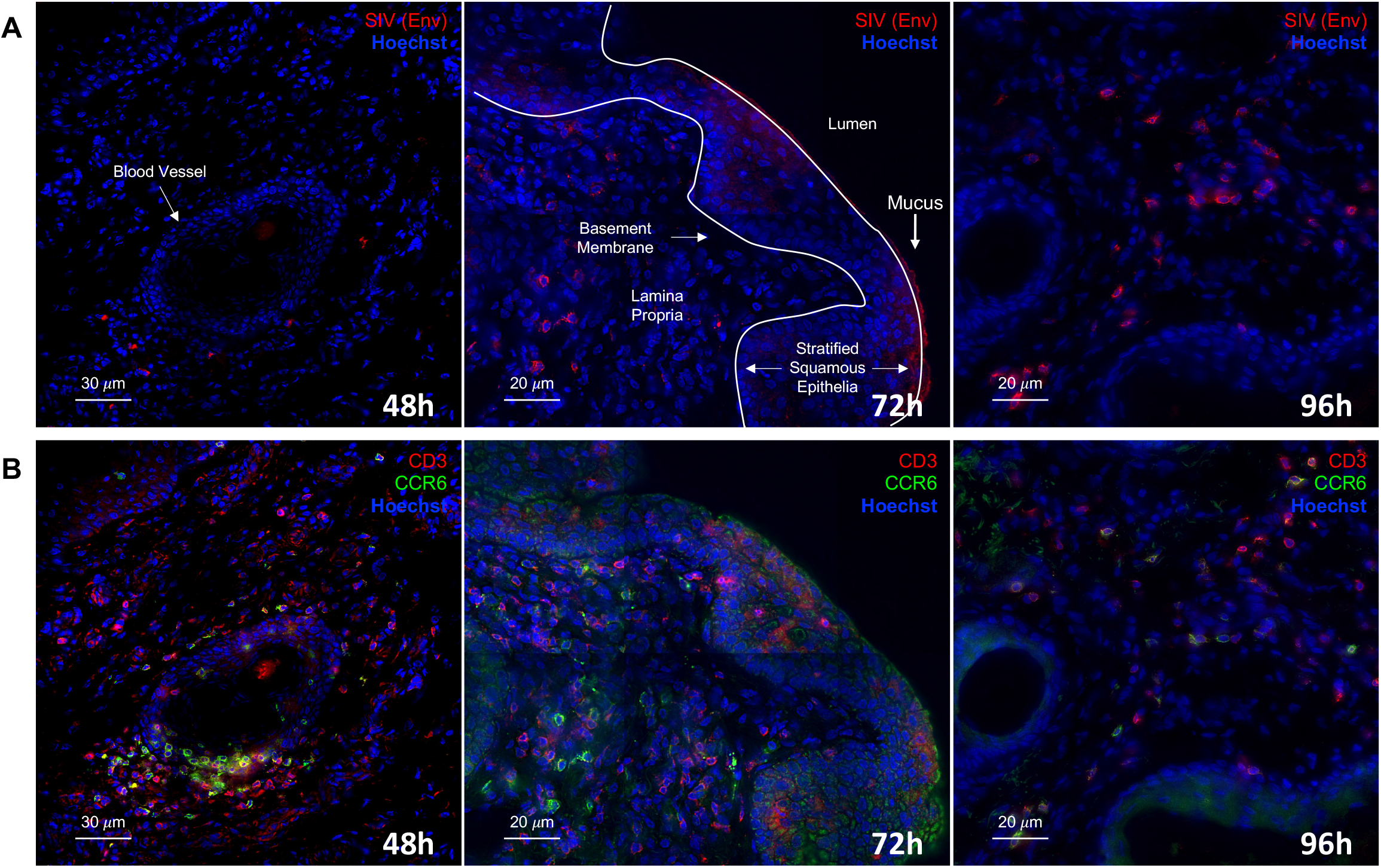
Time dependent changes in the size of infected cell foci and infected cell phenotyping in anal tissue. Cryosections of anal tissue were triple immunolabeled for SIV Env, CD3, and CCR6. Montage of multiple low-magnification (40X) images of immunolabeled and Hoechst stained cryosections depicting a large section of anal tissue at three different time points. The striated nuclear organization reveals a thick stratified epithelial layer of the anal tissue with a layer of autofluorescent mucus atop of it. Arrows point to the other important structural features including the lumen, basement membrane, blood vessel, and lamina propria where the majority of SIV infected (A) and target cells (B) reside.

**Figure 4:**
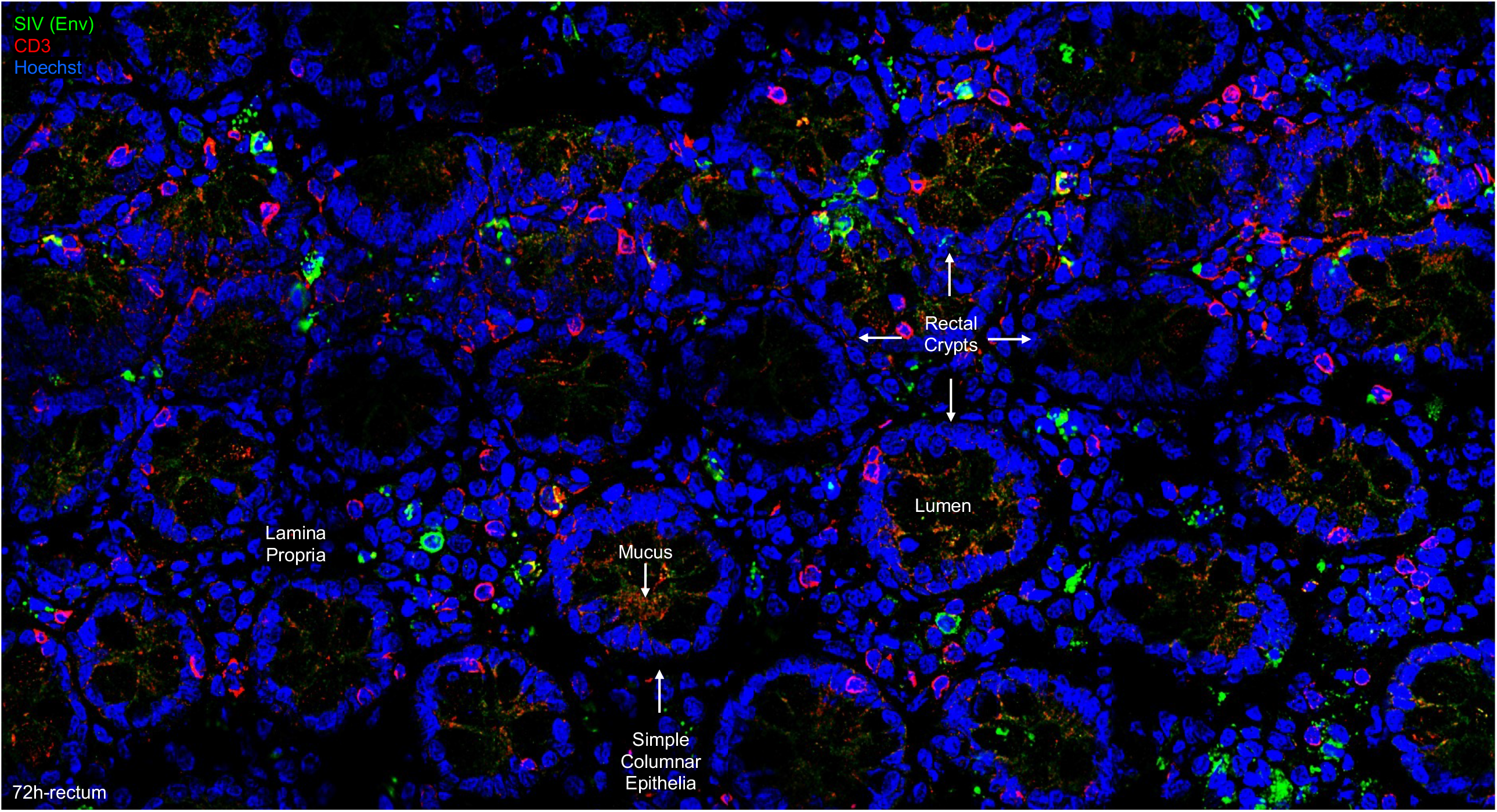
Infected cell distribution and immune phenotyping at the rectal mucosa. Cryosections of rectal tissue were triple immunolabeled for SIV Env, CD3, and CCR6. Montage of multiple low-magnification (40X) images of an immunolabeled and Hoechst stained cryosection depicting a large section of rectal tissue 72h post rectal challenge. The circular organization of nuclei reveals the mucus producing rectal crypts in simple columnar epithelia of these tissues. SIV infected and target cells primarily reside in the lamina propria region while some are also observed within the simple columnar layer of the epithelia.

### Heterogeneous infected cell phenotypes in anorectal tissues are consistent over time

In our previous studies, we used CD3 and CCR6 antibodies to phenotype transduced cells and to group them into one of the four categories including: Th17 T cells (CD3+ CCR6+), other T cells (CD3+ CCR6-), iDCs (CD3-CCR6+) and undefined, myeloid-like cells (CD3-CCR6-). Using the same methodology, we phenotyped a total of 2,377 SIV-infected cells at anorectal mucosa at 48h (n=404), 72h (n=870) and 96h (n=1,103) after rectal inoculation (Figure 5A and Table 1). Phenotyping infected cells at three distinct time points revealed that a wide variety of infected cells including but not limited to Th17 T cells, non Th17 T cells, iDCs and myeloid-like cells represent early targets. When we looked at the proportions of these four cell types in the anus and rectum together, we noted little change over the first 4 days of infection (Figure 5B).

**TABLE 1.** CD3 and CCR6 phenotypic analysis of infected cells at anorectal tissues after anorectal inoculation with SIVmac239 virus

| Animal | Time Point | No. (%) of CD3+ cells that are: |  | No. (%) of CD3+ cells that are: |  | Total no. of infected cells |
| --- | --- | --- | --- | --- | --- | --- |
|  |  | CCR6<sup>+</sup> | CCR6<sup>-</sup> | CCR6<sup>+</sup> | CCR6<sup>-</sup> |  |
| 1_CM38 | 48h | 29 (38) | 0 (0) | 43 (56) | 5 (6) | 77 |
| 2_IN07 | 48h | 45 (23) | 12 (6) | 57 (29) | 81 (42) | 195 |
| 3_EB11 | 48h | 5 (12) | 11 (26) | 6 (14) | 21 (49) | 43 |
| 4_DK09 | 48h | 44 (49) | 7 (8) | 27 (30) | 11 (12) | 89 |
| **Total no. of infected cells (% avg ± SD [%])** |  | **123 (31±16)** | **30 (10±11)** | **133 (32±17 )** | **118 (27±21)** | **404** |
| 5_GK29 | 72h | 85 (66) | 10 (8) | 19 (15) | 15 (12) | 129 |
| 6_KL24 | 72h | 48 (11) | 23 (5) | 165 (39) | 187 (44) | 423 |
| 7_HR73 | 72h | 45 (32) | 26 (19) | 30 (21) | 39 (28) | 140 |
| 8_II82 | 72h | 35 (20) | 19 (11) | 22 (12) | 102 (57) | 178 |
| **Total no. of infected cells (% avg ± SD [%])** |  | **213 (32±24)** | **78 (11±6)** | **236 (22±12)** | **343 (35±19)** | **870** |
| 9_HR28 | 96h | 55 (31) | 43 (24) | 23 (13) | 55 (31) | 176 |
| 10_DN77 | 96h | 175 (33) | 107 (21) | 180 (35) | 57 (11) | 519 |
| 11_JP82 | 96h | 75 (36) | 28 (13) | 85 (41) | 21 (10) | 209 |
| 12_KD09 | 96h | 41 (21) | 26 (13) | 45 (23) | 87 (44) | 199 |
| **Total no. of infected cells (% avg ± SD [%])** |  | **346 (30±6)** | **204 (18±6)** | **333 (28±12)** | **220 (24±16)** | **1103** |

**Figure 5:**
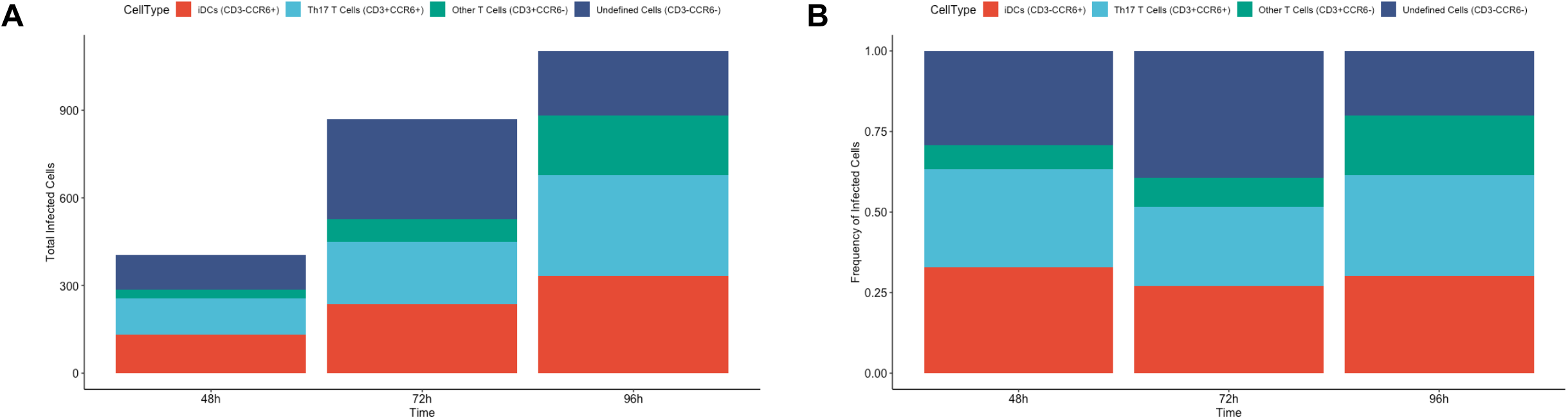
Cell counts and proportions of SIV infected cell types in anorectal tissues. Cryosections of the anorectal tissue were triple immunolabeled for SIV Env, CD3, and CCR6 and infected cells were grouped into one of four categories based on the expression of CD3 and CCR6. (A) Total number of infected cells for each of the four phenotypes at 48h, 72h and 96h time points. (B) Proportion of the total infected cells for each cell type. The four cell phenotypes identified were displayed as a percentage (of 100%) at three different time points.

### Infected cell phenotypes differ in squamous and columnar epithelia

Given that the anus and rectum are composed of tissues that are structurally quite different [32], we decided to look at the same data on a tissue-specific basis. Based on record keeping during tissue dissections and tissue appearance under the microscope, infected cells were either placed in anus or rectum group. The small subset of cells was of unknown tissue origin and were therefore not included in the follow up analysis. Figure 6A and Table 2 show the total number of SIV-infected cells phenotyped with CD3 and CCR6 antibodies at each of the three time points in the anus and rectum separately. We were able to find infected cells in both tissues at all time points, however, the lowest numbers of infected cells were found in the anus tissue 48h after the challenge. Comparing the anus and rectum directly revealed different patterns in infected cell phenotype dynamics throughout infection (Figure 6B). When we tested for statistically significant changes, we found that in the anus the biggest temporal change occurs in infected Th17 T cells at 72hrs and 96hrs (FDR=0.03) and in undefined, myeloid-like, cells at 96hrs (FDR=0.03), both compared to 48h post challenge levels (Figure 7A). In the rectum, however, the biggest temporal change occurs in the infection rate of other T cells, which likewise significantly increased in number both at 72h and 96h compared to 48h (FDR=0.02) as the infection progressed (Figure 7B).

**TABLE 2.** CD3 and CCR6 phenotypic analysis of infected cells anal vs. rectal tissues after anorectal inoculation with SIVmac239 virus

| Animal | Time Point | ANAL |  |  |  |  | RECTAL |  |  |  |  |
| --- | --- | --- | --- | --- | --- | --- | --- | --- | --- | --- | --- |
|  |  | No. (%) of CD3+ cells that are: |  | No. (%) of CD3+ cells that are: |  | Total no. of infected cells | No. (%) of CD3+ cells that are: |  | No. (%) of CD3+ cells that are: |  | Total no. of infected cells |
|  |  | CCR6<sup>+</sup> | CCR6<sup>-</sup> | CCR6<sup>+</sup> | CCR6<sup>-</sup> |  | CCR6<sup>+</sup> | CCR6<sup>-</sup> | CCR6<sup>+</sup> | CCR6<sup>-</sup> |  |
| 1_CM38 | 48h | 0 (0) | 0 (0) | 0 (0) | 0 (0) | 0 | 0 (0) | 0 (0) | 0 (0) | 0 (0) | 0 |
| 2_IN07 | 48h | 3 (60) | 1 (20) | 0 (0) | 1 (20) | 5 | 9 (8) | 7 (6) | 17 (15) | 80 (71) | 113 |
| 3_EB11 | 48h | 0 (0) | 0 (0) | 0 (0) | 0 (0) | 0 | 0 (0) | 0 (0) | 0 (0) | 0 (0) | 0 |
| 4_DK09 | 48h | 3 (30) | 6 (60) | 0 (0) | 1 (10) | 10 | 41 (52) | 1 (1) | 27 (34) | 10 (13) | 79 |
| Total no. of infected cells (% avg ± SD [%]) |  | 6 (45±21) | 7 (40±28) | 0 (0±0) | 2 (15±7) | 15 | 50 (30±31) | 8 (4±4) | 44 (25±13) | 90 (42±41) | 192 |
| 5_GK29 | 72h | 80 (84) | 0 (0) | 15 (16) | 0 (0) | 95 | 5 (15) | 10 (29) | 4 (12) | 15 (44) | 34 |
| 6_KL24 | 72h | 27 (21) | 8 (6) | 83 (65) | 10 (8) | 128 | 21 (7) | 15 (5) | 82 (28) | 177 (60) | 295 |
| 7_HR73 | 72h | 16 (67) | 1 (4) | 5 (21) | 2 (8) | 24 | 29 (25) | 25 (22) | 25 (22) | 37 (32) | 116 |
| 8_II82 | 72h | 0 (0) | 0 (0) | 0 (0) | 0 (0) | 0 | 35 (20) | 19 (11) | 22 (12) | 102 (57) | 178 |
| Total no. of infected cells (% avg ± SD [%]) |  | 123 (57±32) | 9 (3±4) | 103 (34±27) | 12 (5±5) | 247 | 90 (17±8) | 69 (17±11) | 133 (19±8) | 331 (48±13) | 623 |
| 9_HR28 | 96h | 39 (41) | 13 (14) | 14 (15) | 28 (30) | 94 | 0 (0) | 4 (11) | 6 (17) | 25 (71) | 35 |
| 10_DN77 | 96h | 109 (30) | 45 (12) | 173 (47) | 42 (11) | 369 | 27 (35) | 30 (40) | 6 (8) | 13 (17) | 76 |
| 11_JP82 | 96h | 1 (14) | 0 (0) | 1 (14) | 5 (71) | 7 | 69 (38) | 16 (9) | 84 (46) | 14 (8) | 183 |
| 12_KD09 | 96h | 0 (0) | 0 (0) | 0 (0) | 0 (0) | 0 | 41 (21) | 26 (13) | 45 (23) | 87 (44) | 199 |
| Total no. of infected cells (% avg ± SD [%]) |  | 149 (28±14) | 58 (9±8) | 188 (25±19) | 75 (37±31) | 470 | 137 (24±17) | 76 (18±15) | 141 (23±16) | 139 (35±28) | 493 |

**Figure 6:**
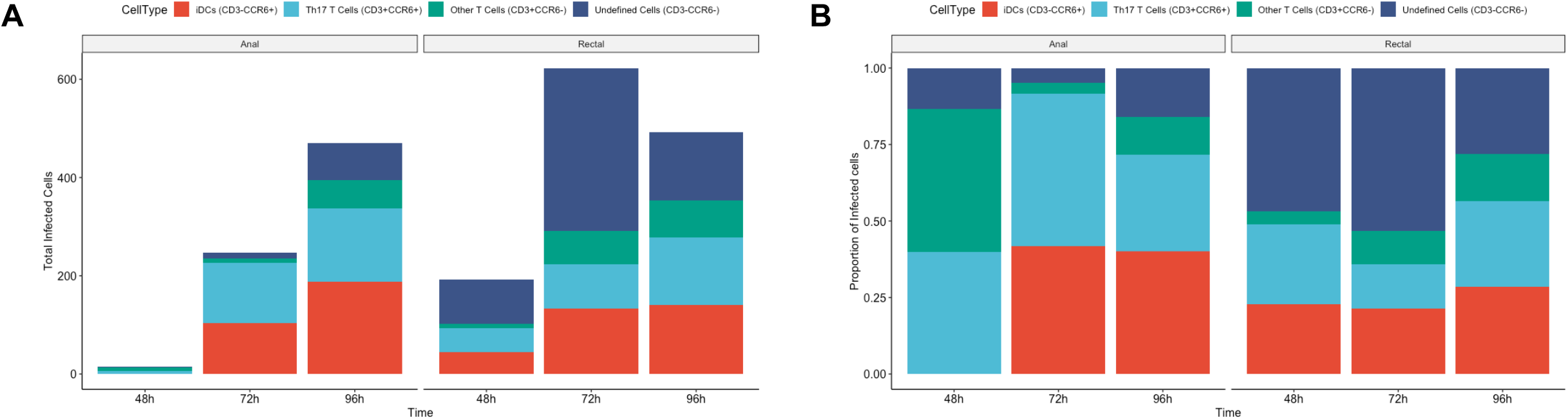
Cell counts and proportions of SIV infected cell types in anus versus rectum. The same SIV-infected cells of four different phenotypes were analyzed in the anus and rectum separately. (A) Total number of infected cells for each of the four phenotypes at 48h, 72h and 96h time points in the anus and rectum separately. (B) Proportion of the total infected cells for each cell type per tissue. The four cell phenotypes identified were displayed as a percentage at three different time points in the anus and rectum separately.

**Figure 7:**
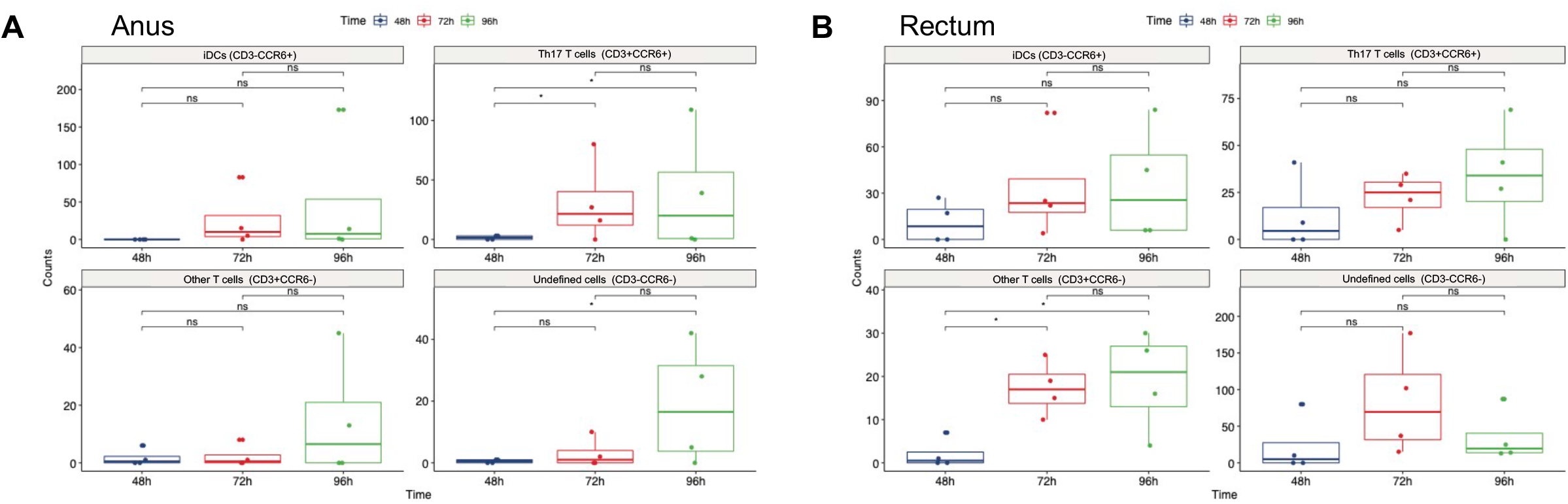
Statistical analysis of temporal changes in infected cell phenotypes in anus and rectum. Horizontal lines in each box represent the median value and the lower and upper error bars are the interquartile ranges. Significance is indicated for the comparisons performed within each fitted model, ns=non-significant, an asterisk indicates significance (FDR<0.05).

## DISCUSSION

The objective of this study was to gain insights into early events that take place in the anorectal mucosa following RAI by utilizing the rhesus macaque rectal challenge model. We were specifically interested in learning which sites are permissive to the virus, elucidating the phenotypes of the infected cells in the first four days of infection and determining the roles different tissue types play in virus susceptibility and progression. By using a mixed challenge inoculum with non-replicative reporter virus and wild-type SIVmac239 we were able to identify sites that are permissive to the virus and consistently identify infected cells on days two, three and four post rectal challenge. The susceptibility of both anal and rectal tissues was confirmed by immunofluorescence microscopy as small regions of tissue expressing luciferase also contained infected cells identified by staining for SIV proteins Env and Gag (Figure 1B). The permissiveness of anus tissue is noteworthy given that previous efforts to develop vaccines and microbicides have been mostly focused on rectal tissues which are historically thought to be more permissive due to their single columnar epithelial nature and richness in diverse populations of HIV-1 target cells.

A high density of HIV-1 target cells ranging in phenotypes was notable in both anal and rectal tissues (Figure 3B and Figure 4) in agreement with the study by McElrath, et al. which demonstrated that human gut tissues contain diverse populations of HIV target cells. This study also found greater CCR5 expression on rectal macrophages in comparison to distal gut macrophages, suggesting increased HIV susceptibility towards the anus [33]. The infected cells we identified were found in the areas rich in immune cells and they mostly localized to the lamina propria, proximal to the basement membrane (Figure 3A and Figure 4). The infected cell had a focal appearance and increased in size as infection progressed; 48h after the challenge foci consisted of only a few of infected cells, however by 96h post challenge the infected cell foci consisted of hundreds of cells and often stretched over several adjacent tissue sections (Figure 3A).

The subsequent characterization of infected cells revealed that they had varying degrees of CD4 expression; while some infected cells had robust CD4 signals others had little to no CD4 detectable, consistent with previously published work reporting that wild-type viruses can cause down-modulation of CD4 cell surface expression via Nef protein in productively infected cells [34]. We also observed varying levels of T cell specific markers in some of the infected cells and found that both T cells and non-T cells are susceptible to the virus as early as 48h after rectal challenge. Positivity for the viral proteins Env and Gag along with expected changes in CD4 and CD3 expression indicated we are successfully detecting the first infected T cells and non-T cells at the anorectal mucosa of rectally challenged macaques.

Our previous work demonstrated that the vast majority of CD3+CCR6+ cells and all CD3-CCR6+ cells at anorectal mucosa express RORγT, a Th17 lineage specific factor and DC-specific intercellular adhesion molecule-3-grabbing nonintegrin (DC-SIGN) molecule, respectively [35, 36], thus allowing us to group the infected cells into one of the four major categories including Th17 T cells, other T cells, iDCs, and myeloid-like, currently uncharacterized, cells based on overall morphology and expression or lack of these two markers.

Comparing the proportions of transduced cells (Li670 only challenge) in our original study [22] and SIVmac239 infected cells identified here at the same 48h time point revealed that the wild-type virus infects a much more heterogeneous population of cells in the early course of infection, likely due to the activation of immune responses near or at the site of infection. The transduction events by the Li670 vector were primarily dominated by Th17 T cells which accounted for over 60% of the first cells transduced followed by iDCs at 20% [22]. However, phenotypic profiling of SIV-infected cells presented here shows comparable ratios of infected Th17 T cells, iDC, and myeloid-like cells at the same time point when tissues of anus and rectum were analyzed together (Figure 5B). Both the Li670 vector and the wild-type virus targeted low numbers of other T cells in the early course of infection (48h) [22] and (Figure 5B). As the infection progressed from two days to four days post challenge, the ratios of infected cell types remained relatively constant when infected cells from the anus and rectum were analyzed together. Given that squamous tissues of anus and columnar tissues of rectum are structurally distinct we set to examine the acquired data in each tissue separately. Based on the tissue morphology, infected cells that were identified in this study were either placed in anus or rectum category. A small subset of cells was of unknown tissue origin and was therefore not included in the follow up analysis. Analyzing the results on a tissue-specific basis revealed that the infected cell phenotypes in anus and rectum are present at different proportions and have distinct temporal dynamics (Figure 6B). In the anus, the biggest temporal change occurred in infected Th17 T cells and myeloid-like cells, which both increased significantly from day 2 to day 4 post challenge (Figure 7A), and for the rectum, the biggest temporal change was notable in the other T cells, which significantly increased from 48h to 72h as well as from 48h to 96h post challenge (Figure 7B). These data highlight the distinct but equally important roles the different tissues play in virus susceptibility and progression.

Lastly, we hoped to be able to compare the proportions of infected cells in anorectal vs. vaginal challenge as our previously published study utilized a similar mixed inoculum in the rhesus macaque vaginal challenge [27]. Given that FRT samples included in the study almost exclusively consist of tissues with stratified squamous epithelia (such as labia, vagina and ectocervix) we sought to compare data from the published FRT study to the anus data presented here. Unfortunately, the number of SIV-infected cells in the anus at 48h time point is very low and (Figure 6A) hence we were not able to make any meaningful comparisons.

While informative this study also has some limitations. For one, imaging tissue sections is not ideal for the analysis of complex biological processes which are three dimensional phenomena. When imaging large foci at 96h time point it is impossible to discern where one focus of infection ends and where a new one begins. To learn more about how foci of infection are formed and how they evolve over the time, our follow up work will rely on using alternative methods such as tissue clearing and light sheet imaging to gain a three-dimensional view of virus dissemination in the intact tissues. Also, Luciferase-IVIS assay is unable to detect more diffuse signals as well as those that are farther away from the surface of the tissue. Therefore, in our follow up work, we will be using the immune PET/CT method to look for more isolated events. Lastly, this study focused on the anus and distal six cm of the adjoining rectal tissue, and therefore did not examine the infection events in more proximal gastrointestinal (GI) tissue which could have different infection dynamics. Work from our own laboratory involving infant rhesus macaques in fact demonstrated that the upper gastrointestinal tract contains the largest number of infected cells following oral challenge [37]. Whole animal imaging with PET/CT will be very informative in understanding infection dynamics in the proximal GI tract.

Collectively, these studies point to the importance of controlling HIV infection in anal tissue in addition to rectal tissue and reveal important distinctions in these two tissues when it comes to early viral dynamics after rectal challenge. The rhesus macaque model utilizing a mixed inoculum challenge is well suited for the investigation of the early events in HIV-1 transmission during RAI and has the potential to inform future development of effective prevention strategies.

## MATERIALS AND METHODS

### Ethics statement

All animal studies were conducted in accordance with protocol P0153, which was reviewed and approved by both Northwestern University and Tulane National Primate Research Center Institutional Animal Care and Use Committees (IACUC). Animals in this study were euthanized by sedation with ketamine-hydrochloride injection followed by intravenous barbiturate overdose, in accordance with the recommendations of the Panel on Euthanasia of the American Veterinary Medical Association.

### Reporter vector and wild-type virus

The LI670-JRFL reporter virus developed in our original study [22] was mixed in inoculum with wild-type SIVmac239 [27] to rectally inoculate twelve female rhesus macaques. In summary, LI670-JRFL reporter was produced by transfecting 293T cells with the following plasmids: 1, 2, 3, and 4. Viruses containing supernatants were collected 48h post-transfection, purified through 0.22 mm filters, concentrated via sucrose gradient and resuspended in DMEM. The concentrated virus was stored at −80 degrees C until use. SIVmac239 was generated by transfection and expansion in CEMx174 cells. Virus production was monitored by measuring the p27 antigen. Cells were maintained with RPMI and the addition of new cells every 3–4 days. At 24h before harvest, cells are washed and resuspended in fresh RPMI. Virus-containing supernatant is then centrifuged, filtered, aliquoted, and stored at –80°C. The p27 concentration of the challenge stock was 1,130 pgmL−1. Infection of Ghost-Bonzo cells revealed infection of ∼4% of the cells at 1:2 dilution 48 hr after viral exposure. Infectivity of viral stock was determined by measuring infection of CEMx174 cells at the limiting viral dilution that was positive for p27 after 21 days of culture; this gave a TCID50 determination of 105.91 per ml.

### Non-human primate studies

All animal handling was carried out at the Tulane National Primate Research Center. This study involved twelve female rhesus macaques which were evenly divided into three experimental groups. Prior to challenge, each animal received four to six rectal biopsies at random and was inoculated rectally with a mixture of LI670-VSVg virus and wild-type SIVmac239. Animals were sacrificed either 48h, 72h or 96h after the rectal inoculation (four animals per group). After the designated time animals were euthanized and necropsied. Anorectal tissues encompassing the first 4-6cm of the distal colon were excised in one piece for the purpose of this study and were shipped overnight in RPMI medium on ice for further processing and analysis by the Hope Laboratory at Northwestern University.

### *In vivo* imaging system

Following the arrival of tissues at Northwestern University anorectal tissues were removed from the RPMI medium and rinsed several times in cold phosphate-buffered saline (PBS) to remove any debris and fecal matter. The adjoining anus and rectum were imaged in one piece in the PerkinElmer IVIS Lumina Series III device to capture background luminescence. Tissues were then soaked in 100 mM D-Luciferin (Biosynth) for ten minutes, and re-imaged with IVIS under the same exposure settings. Regions of tissue with robust luciferase expression were noted and all tissue pieces were frozen in optimal cutting temperature (OCT) medium as described previously [22, 27]. Attention was taken to demark whether the tissue was from the anus or rectum.

### Immunolabeling and microscopy

Frozen tissue blocks of the anus and rectum containing high luciferase expression were cryosectioned at 10-20um thickness and placed on a glass slide. Thin tissue sections were fixed with 1% formaldehyde in PIPES [piperazine-N, N’-bis(2-ethanesulfonic acid)] buffer (Sigma) for ten minutes at room temperature and blocked with cold normal donkey serum (Jackson ImmunoResearch) for thirty minutes at room temperature. Primary antibodies used in this study included: SIV Gag (Ag3.0, hybridoma supernatant), SIV Env (KK41 and KK42, AIDS Reagent Repository), CD3 (SP7, Abcam), CD4 (OKT4, hybridoma supernatant) and CCR6 (53103, R&D Systems). All secondary antibodies were raised in donkeys, were diluted at 1:500, and were purchased from Jackson ImmunoResearch. Normal donkey serum was used as a dilutant for all primary and secondary antibodies in this study. Primary and secondary antibodies were incubated for 1 hour or 30 minutes, respectively at ambient temperature. When two primary antibodies of the same species were used, one was labeled with a Zenon IgG labeling kit (ThermoFisher Scientific) using AlexaFluor fluorochromes, according to the manufacturer’s instructions. After antibody incubations nuclear stain Hoechst (Thermo Fisher Scientific, 1:25,000 dilution in cold PBS) was applied for five minutes at room temperature. Secondary-only stained slides were used to set up the threshold for each channel when setting up the experiments. Image stacks (20-40 sections in Z-plane in 0.5-um steps) were acquired by DeltaVision Elite (GE) inverted microscope equipped with 40- and 60x magnification objectives and deconvolved using SoftWoRx software.

### Statistical analysis

The data from the two tissues were separated and fitted to a Negative Binomial generalized linear model to test for temporal differences in cell counts per cell type. We included time point, cell type, and their interaction in the model and tested for changes in cell types per time using Benjamini-Hochberg method for multiple testing adjustment. FDR was set at FDR < 0.05 for significance.

## ACKNOWLEDGEMENTS AND AUTHORS’ CONTRIBUTIONS

We thank Edgar Matias and Sixia Xiao for help with tissue sectioning, Edward Allen for virus production, Gianguido Cianci for valuable discussions and Elena Martinelli and Eileen Porter for editing the manuscript.

We have no conflicts of interest to declare.

DM performed tissue processing, conducted experiments, analyzed the data, generated figures, and wrote the manuscript. LC and NG performed experiments, analyzed the data, and generated figures. RRL performed statistical analysis and generated figures. MDM coordinated animal challenges and helped with tissue processing. RSV conducted animal challenges and supervised animal necropsies. TJH designed the study and provided feedback at all stages from experimental planning to manuscript editing. All authors were given a chance to read the manuscript and provide feedback during the editing process.

This work was funded by National Institutes of Health grants 1UM1AI120184-01 and R37AI094595 and supplement to R01AI94595 awarded to TJH.

## BIBLIOGRAPHY

1. Elmes, J., et al., Receptive anal sex contributes substantially to heterosexually acquired HIV infections among at-risk women in twenty US cities: Results from a modelling analysis. Am J Reprod Immunol, 2020. 84(2): p. e13263.

2. Baggaley, R.F., R.G. White, and M.C. Boily, HIV transmission risk through anal intercourse: systematic review, meta-analysis and implications for HIV prevention. Int J Epidemiol, 2010. 39(4): p. 1048–63.

3. Baggaley, R.F., et al., Does per-act HIV-1 transmission risk through anal sex vary by gender? An updated systematic review and meta-analysis. Am J Reprod Immunol, 2018. 80(5): p. e13039.

4. Biello, K.B., et al., MSM at Highest Risk for HIV Acquisition Express Greatest Interest and Preference for Injectable Antiretroviral PrEP Compared to Daily, Oral Medication. AIDS Behav, 2018. 22(4): p. 1158–1164.

5. Kelley, C.F., et al., The rectal mucosa and condomless receptive anal intercourse in HIV-negative MSM: implications for HIV transmission and prevention. Mucosal Immunol, 2017. 10(4): p. 996–1007.

6. Ribeiro Dos Santos, P., et al., Rapid dissemination of SIV follows multisite entry after rectal inoculation. PLoS One, 2011. 6(5): p. e19493.

7. Hladik, F. and M.J. McElrath, Setting the stage: host invasion by HIV. Nat Rev Immunol, 2008. 8(6): p. 447–57.

8. Haase, A.T., Targeting early infection to prevent HIV-1 mucosal transmission. Nature, 2010. 464(7286): p. 217–23.

9. Keele, B.F. and J.D. Estes, Barriers to mucosal transmission of immunodeficiency viruses. Blood, 2011. 118(4): p. 839–46.

10. Gupta, P., et al., Memory CD4(+) T cells are the earliest detectable human immunodeficiency virus type 1 (HIV-1)-infected cells in the female genital mucosal tissue during HIV-1 transmission in an organ culture system. J Virol, 2002. 76(19): p. 9868–76.

11. Rodriguez-Garcia, M., et al., Dendritic cells from the human female reproductive tract rapidly capture and respond to HIV. Mucosal Immunol, 2017. 10(2): p. 531–544.

12. Mattei, D., et al., Enzymatic Dissociation Induces Transcriptional and Proteotype Bias in Brain Cell Populations. Int J Mol Sci, 2020. 21(21).

13. Tighe, R.M., et al., Improving the Quality and Reproducibility of Flow Cytometry in the Lung. An Official American Thoracic Society Workshop Report. Am J Respir Cell Mol Biol, 2019. 61(2): p. 150–161.

14. Taghizadeh, R.R., K.J. Cetrulo, and C.L. Cetrulo, Collagenase Impacts the Quantity and Quality of Native Mesenchymal Stem/Stromal Cells Derived during Processing of Umbilical Cord Tissue. Cell Transplant, 2018. 27(1): p. 181–193.

15. Sattentau, Q.J. and M. Stevenson, Macrophages and HIV-1: An Unhealthy Constellation. Cell Host Microbe, 2016. 19(3): p. 304–10.

16. Geijtenbeek, T.B., et al., DC-SIGN, a dendritic cell-specific HIV-1-binding protein that enhances trans-infection of T cells. Cell, 2000. 100(5): p. 587–97.

17. Cory, T.J., et al., Overcoming pharmacologic sanctuaries. Curr Opin HIV AIDS, 2013. 8(3): p. 190–5.

18. Stieh, D.J., et al., Vaginal challenge with an SIV-based dual reporter system reveals that infection can occur throughout the upper and lower female reproductive tract. PLoS Pathog, 2014. 10(10): p. e1004440.

19. Rabinovich, B.A., et al., Visualizing fewer than 10 mouse T cells with an enhanced firefly luciferase in immunocompetent mouse models of cancer. Proc Natl Acad Sci U S A, 2008. 105(38): p. 14342–6.

20. Shcherbakova, D.M. and V.V. Verkhusha, Near-infrared fluorescent proteins for multicolor in vivo imaging. Nat Methods, 2013. 10(8): p. 751–4.

21. Jun, Y.W., et al., Addressing the autofluorescence issue in deep tissue imaging by two-photon microscopy: the significance of far-red emitting dyes. Chem Sci, 2017. 8(11): p. 7696–7704.

22. Maric, D., et al., Th17 T Cells and Immature Dendritic Cells Are the Preferential Initial Targets after Rectal Challenge with a Simian Immunodeficiency Virus-Based Replication-Defective Dual-Reporter Vector. J Virol, 2021. 95(19): p. e0070721.

23. Rodriguez-Garcia, M., et al., Phenotype and susceptibility to HIV infection of CD4+ Th17 cells in the human female reproductive tract. Mucosal Immunol, 2014. 7(6): p. 1375–85.

24. Alvarez, Y., et al., Preferential HIV infection of CCR6+ Th17 cells is associated with higher levels of virus receptor expression and lack of CCR5 ligands. J Virol, 2013. 87(19): p. 10843–54.

25. Yang, X.O., et al., T helper 17 lineage differentiation is programmed by orphan nuclear receptors ROR alpha and ROR gamma. Immunity, 2008. 28(1): p. 29–39.

26. Wu, Z., T. Hu, and P. Kaiser, Chicken CCR6 and CCR7 are markers for immature and mature dendritic cells respectively. Dev Comp Immunol, 2011. 35(5): p. 563–7.

27. Stieh, D.J., et al., Th17 Cells Are Preferentially Infected Very Early after Vaginal Transmission of SIV in Macaques. Cell Host Microbe, 2016. 19(4): p. 529–40.

28. Bour, S., R. Geleziunas, and M.A. Wainberg, The human immunodeficiency virus type 1 (HIV-1) CD4 receptor and its central role in promotion of HIV-1 infection. Microbiol Rev, 1995. 59(1): p. 63–93.

29. Woolley, D.W. and B.W. Gommi, Serotonin Receptors. Iv. Specific Deficiency of Receptors in Galactose Poisoning and Its Possible Relationship to the Idiocy of Galactosemia. Proc Natl Acad Sci U S A, 1964. 52: p. 14–9.

30. Yang, H., R.M. Parkhouse, and T. Wileman, Monoclonal antibodies that identify the CD3 molecules expressed specifically at the surface of porcine gammadelta-T cells. Immunology, 2005. 115(2): p. 189–96.

31. Clevers, H., et al., The T cell receptor/CD3 complex: a dynamic protein ensemble. Annu Rev Immunol, 1988. 6: p. 629–62.

32. Tanaka, E., et al., Morphology of the epithelium of the lower rectum and the anal canal in the adult human. Med Mol Morphol, 2012. 45(2): p. 72–9.

33. McElrath, M.J., et al., Comprehensive assessment of HIV target cells in the distal human gut suggests increasing HIV susceptibility toward the anus. J Acquir Immune Defic Syndr, 2013. 63(3): p. 263–71.

34. Garcia, J.V. and A.D. Miller, Serine phosphorylation-independent downregulation of cell-surface CD4 by nef. Nature, 1991. 350(6318): p. 508–11.

35. Granelli-Piperno, A., et al., Dendritic cell-specific intercellular adhesion molecule 3-grabbing nonintegrin/CD209 is abundant on macrophages in the normal human lymph node and is not required for dendritic cell stimulation of the mixed leukocyte reaction. J Immunol, 2005. 175(7): p. 4265–73.

36. Ruan, Q., et al., The Th17 immune response is controlled by the Rel-RORgamma-RORgamma T transcriptional axis. J Exp Med, 2011. 208(11): p. 2321–33.

37. Taylor, R.A., et al., Localization of infection in neonatal rhesus macaques after oral viral challenge. PLoS Pathog, 2021. 17(11): p. e1009855.

